# A Lightweight, High-Throughput Classifier for North American Insects Using EfficientNet: Elytra 1.0

**DOI:** 10.64898/2026.02.16.706225

**Authors:** Nicholas Aflitto

## Abstract

Large-scale biodiversity monitoring is often inhibited by taxonomic obstacles. While deep learning has demonstrated efficacy in species identification, the increasing reliance on large Vision Transformers (ViTs) creates computational barriers that restrict usage to cloud-based infrastructure. Recent foundation models, such as BioCLIP and the Insect-1M framework, require parameter counts exceeding 100M, rendering them unsuitable for edge deployment in field operations. This study presents Elytra 1.0, a computer vision model optimized for edge deployment and capable of classifying 3,127 common North American insect species. The dataset, comprising 2.6 million images, includes all insect species in North America with over 1,000 research-grade observations on iNaturalist. An EfficientNet-B0 architecture was trained using transfer learning from ImageNet with adaptive learning rate scheduling. The model achieved 91.27% Top-1 Accuracy and 97.6% Top-5 Accuracy on an internal test set (N=289,151 images). To rigorously evaluate generalization beyond photographer-specific patterns, an independent observer-excluded test set (N=5,780 images, 578 species) was constructed comprising images exclusively from photographers who contributed zero training data. A post-hoc spatiotemporal audit revealed this test set was heavily skewed toward the Neotropics (Mean Lat: 6.05° N) during the boreal winter (Dec 2025–early Feb 2026). Despite this significant biogeographic and phenological shift from the predominantly temperate training data, the model achieved 86.68% Top-1 Accuracy (95% CI: 85.8–87.5%). This confirms that Elytra 1.0 relies on robust morphological features rather than learning background environmental correlations, maintaining high performance even in novel ecological contexts.The resulting model file size is 30 MB with an inference speed exceeding 700 frames per second (FPS) on mobile hardware. These results indicate that optimized convolutional architectures can achieve competitive accuracy with server-grade transformers while remaining suitable for decentralized, offline monitoring applications.

## 1 Introduction

Insects comprise the majority of terrestrial animal diversity and provide vital ecosystem services, yet they face major global declines (Wagner et al., 2021). Monitoring these trends requires data scales exceeding human capacity, driving the adoption of automated identification tools (Engel et al., 2021). While recent models like InsectNet (Chiranjeevi et al., 2025) demonstrate that deep learning can distinguish tens of thousands of species with >96% accuracy, these systems often rely on global-to-local cloud architectures that are inaccessible in remote environments.

A trade-off has emerged in computational ecology. The field has increasingly adopted massive Vision Transformers (ViTs) (Liu et al., 2022) and Neural Architecture Search (NAS) frameworks like BioAutoML-NAS (Abian et al., 2025). While these models set new accuracy benchmarks, often exceeding 96% on datasets like BIOSCAN-5M, they frequently exceed 500 MB in size and require substantial GPU resources. This limits utility in remote field stations and mobile applications where bandwidth is constrained (Lee & Di Giacomo, 2024).

There is a need for edge AI in entomology, and ecology more broadly, with models that balance taxonomic breadth and computational efficiency (Maslej et al., 2025). Beyond practical deployment constraints, the environmental cost of training and deploying large-scale models has become an important consideration. Recent studies estimate that training a single large vision model can consume as much energy as several households use in a year (Strubell et al., 2019), making lightweight architectures not only more accessible but also more sustainable for long-term biodiversity monitoring efforts. To address this, I developed Elytra 1.0, a classifier optimized for common North American insect species.

## 2 Methods

### 2.1 Dataset Collection and Curation

Training data were sourced from the iNaturalist open dataset (Van Horn et al., 2018), filtered for observations in North America with open licenses (CC0, CC-BY, CC-BY-NC). To help ensure data quality, an inclusion threshold was applied where species were included only if they possessed >1,000 research-grade observations in North America as of December 2025. Research-grade is defined by iNaturalist as observations with a date, location, photo, and community consensus on the ID (2+ agreeing identifications).

The final dataset comprised 3,127 species (2,602,535 images). Unlike typical natural world datasets which suffer from extreme class imbalance, this dataset is explicitly curated for uniformity, with a median of 900 images per class.

Dataset Partitioning and Observer Diversity: The dataset was partitioned into Training (80%, N=2,313,251), Validation (10%, N=289,284), and Internal Test (10%, N=289,151) subsets. The validation subset was used exclusively for hyperparameter tuning and early stopping during model development. The internal test subset was sequestered prior to training and accessed only for final performance evaluation. To assess photographer diversity in the training data, 50 species were sampled and observer IDs were extracted from training images via the iNaturalist API. This audit revealed high photographer diversity (observer diversity score: 1.21), indicating that most training images originated from unique photographers, substantially reducing potential bias from repeated observer patterns.

Observer-Independent Test Set: To rigorously evaluate generalization beyond photographer-specific patterns, an independent test set was constructed that explicitly excluded all photographers who contributed to training data. For each of the 3,127 species, observer IDs were identified from a random sample of training images (N≈30 per species) via the iNaturalist API. iNaturalist was then queried for research-grade observations and up to 10 test images per species were downloaded from photographers not present in the training set. This procedure yielded a test set of 5,780 images spanning 578 species (18.5% of taxonomic range), ensuring complete observer independence.

Spatiotemporal Audit of Test Data: To characterize the distribution of the observer-independent test set relative to the training data, a metadata audit was performed on a random subset of test images (N=200). Observations were queried via the iNaturalist API to extract coordinate and timestamp data. This audit revealed a distinct spatiotemporal shift: 92% of test images were observed during the North American winter (December 2025–January 2026). Consequently, the test set distribution was geographically skewed toward the Neotropical extent of the target species’ ranges (Mean Latitude: 6.05° N; range: Central America to Northern South America). This incidental shift provided a rigorous stress test for the model, evaluating its ability to generalize to tropical background environments and overwintering insect phenotypes not heavily represented in the temperate-dominant training set.

### 2.2 Model Architecture

The EfficientNet-B0 (Tan & Le, 2019) architecture was selected over other lightweight candidates (e.g., MobileNetV3, ResNet-18) due to its superior accuracy-to-parameter ratio. While MobileNetV3 offers marginally faster inference, EfficientNet’s compound scaling facilitates deeper feature extraction important for fine-grained classification, all while remaining within the thermal and memory constraints of consumer hardware. Unlike the recent Swin-AARNet (Xiao et al., 2025), which employs computationally expensive attention mechanisms for pest recognition, EfficientNet-B0 prioritizes low-FLOP efficiency. The network was initialized with weights pre-trained on ImageNet (Deng et al., 2009), and the final classification head was replaced with a linear layer containing 3,127 outputs.

### 2.3 Training Protocol

Training was conducted using PyTorch (Paszke et al., 2019) on Apple Silicon (M1 Ultra, Mac Studio). Images were resized to 224×224 pixels and normalized using standard ImageNet statistics. Standard CrossEntropyLoss was applied. The optimization strategy utilized the Adam algorithm (Kingma & Ba, 2014) with an initial learning rate of 1×10−3 and a batch size of 64. A ReduceLROnPlateau scheduler (patience=2, factor=0.1) dynamically reduced the learning rate when validation accuracy plateaued, eventually reaching 1×10−5. Early stopping (patience=5) terminated training at optimal convergence. This strategy effectively created a two-phase learning regime: an initial high-learning-rate phase for rapid feature discovery, followed by fine-grained refinement at lower learning rates. Hyperparameters are summarized in Table 1.

**Table 1.** Hyperparameter Configuration.

| Parameter | Value |
| --- | --- |
| Architecture | EfficientNet-B0 |
| Input Size | 224x224 |
| Batch Size | 64 |
| Optimizer | Adam |
| Learning Rate | $1e-3 \rightarrow 1e-5$ |
| Scheduler | ReduceLROnPlateau |
| Criterion | CrossEntropyLoss |

### 2.4 Data Augmentation

A robust augmentation pipeline was implemented to simulate field conditions:

- Random Resized Crop: Scale 0.8–1.0.
- Geometric: Random Horizontal Flips (p=0.5) and Rotation (±15°).
- Photometric: Color Jitter (Brightness, Contrast, Saturation; factor=0.2).

### 2.5 Validation and Metrics

Performance was evaluated on both the internal test set (N=289,151) and the observer-independent test set (N=5,780, 578 species). Top-1 and Top-5 accuracy were calculated as the proportion of correct predictions across all test samples. Confidence intervals (95% CI) were computed using the Wilson score interval method.

## 3 Results

### 3.1 Classification Performance

Elytra 1.0 achieved a Top-1 Accuracy of 91.27% and a Top-5 Accuracy of 97.6% (Table 2). When macro-averaged across species (i.e., unweighted mean of per-species accuracies), the model achieved 90.23%, indicating relatively balanced performance across common and rare taxa (Figure 1). The generalization gap between training and test accuracy was minimal, validating the efficacy of the augmentation strategies.

**Table 2.** Elytra 1.0 Performance.

| Metric | Internal Test Set | Observer-Independent Test |
| --- | --- | --- |
| **Dataset Size** | 289,151 images, 3,127 species | 5,780 images, 578 species |
| **Top-1 Accuracy** | 91.27% (95% CI: 91.2–91.3%) | 86.68% (95% CI: 85.8–87.5%) |
| **Top-5 Accuracy** | 97.62% |  |
| **Inference Speed (mobile)** | >700 FPS |  |
| **Model Size (Core ML)** | 30 MB |  |

**Figure 1.**
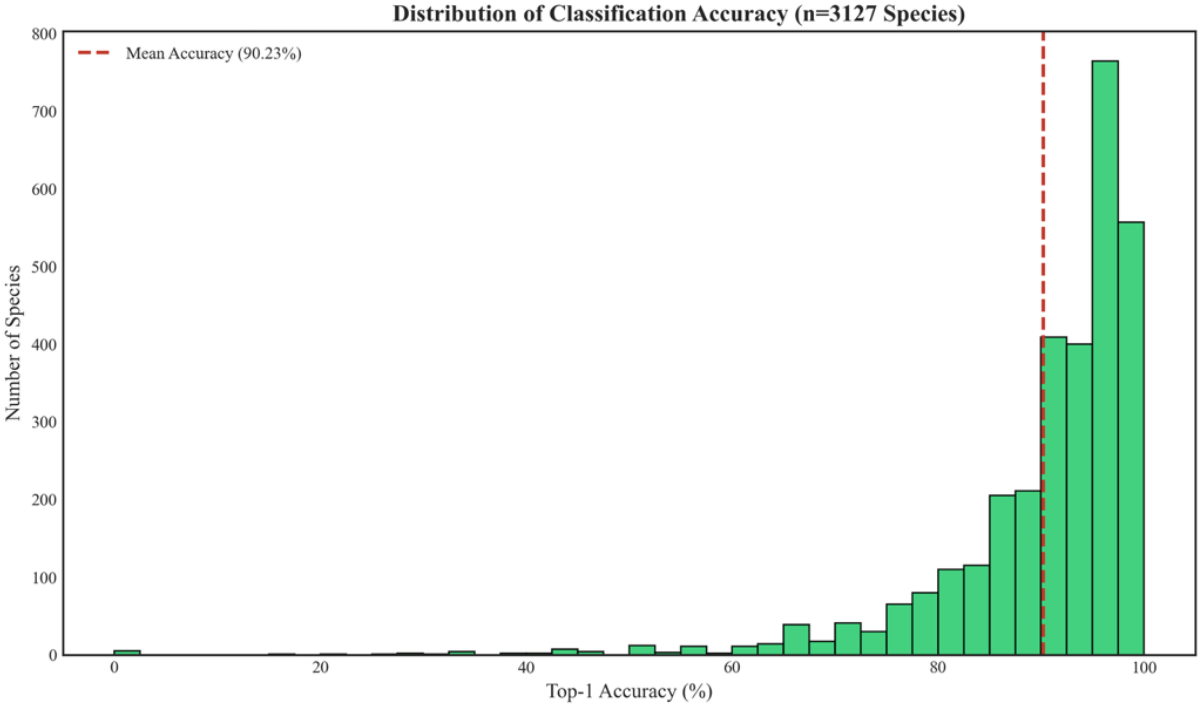
Distribution of per-species classification accuracy across 3,127 taxa. Histogram showing Top-1 accuracy for individual species (n=3,127). The left-skewed distribution (median=92.1%, IQR=87.3–95.4%) indicates that high accuracy is achieved for the majority of species, with only 8.3% of taxa falling below 80% accuracy. Species with <70% accuracy (n=89, 2.8%) are primarily cryptic complexes requiring microscopic features or molecular identification (see Discussion).

To evaluate generalization beyond photographer-specific patterns (observer-independent test performance), the model was tested on 5,780 images from 578 species (18.5% of taxonomic range), comprising exclusively observations from photographers who contributed zero training data. The model achieved 86.68% Top-1 Accuracy (95% CI: 85.8–87.5%) on this observer-independent test set. While this represents a 4.59 percentage point decrease from the internal test set (91.27%), this gap must be interpreted in the context of the test set’s extreme biogeographic shift. The test images predominantly originated from Neotropical winter environments (Lat ∼6° N), presenting lighting conditions, floral backgrounds, and insect life stages (e.g., overwintering adults) distinct from the temperate summer distribution that dominates the training data. The ability to maintain >86% accuracy despite this domain shift indicates strong feature robustness.

Performance varied significantly by taxonomic order (Figure 2). Distinctive groups like Diptera (flies) and Odonata (dragonflies) achieved the highest accuracy, exceeding 92%. Lepidoptera (moths/butterflies) achieved 88.8%, while Coleoptera (beetles) reached 85.1%. In contrast, Hymenoptera (bees/ants/wasps) showed the lowest performance (79.1%), attributable to the high prevalence of cryptic species complexes (e.g., *Lasioglossum spp*.) and Müllerian mimicry rings which often require microscopic examination for reliable identification (Figure 3).

**Figure 2.**
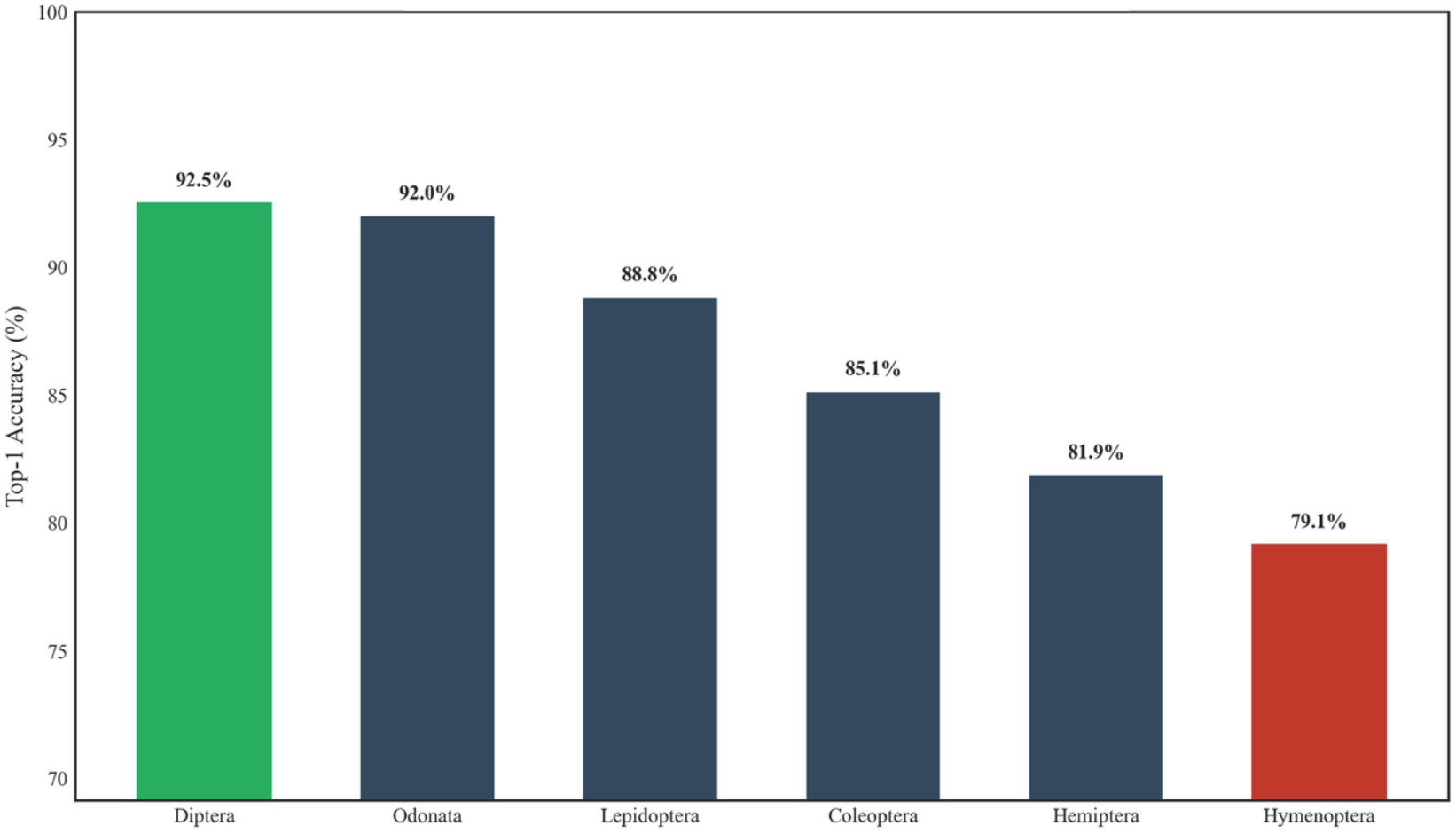
Classification accuracy stratified by taxonomic order. Top-1 accuracy on the internal test set (n=289,151) was highest for Diptera (92.5%) and Odonata (92.0%), likely due to their distinct morphological traits. Hymenoptera achieved the lowest accuracy (79.1%), attributable to the prevalence of cryptic species complexes and Müllerian mimicry rings that complicate visual identification.

**Figure 3.**
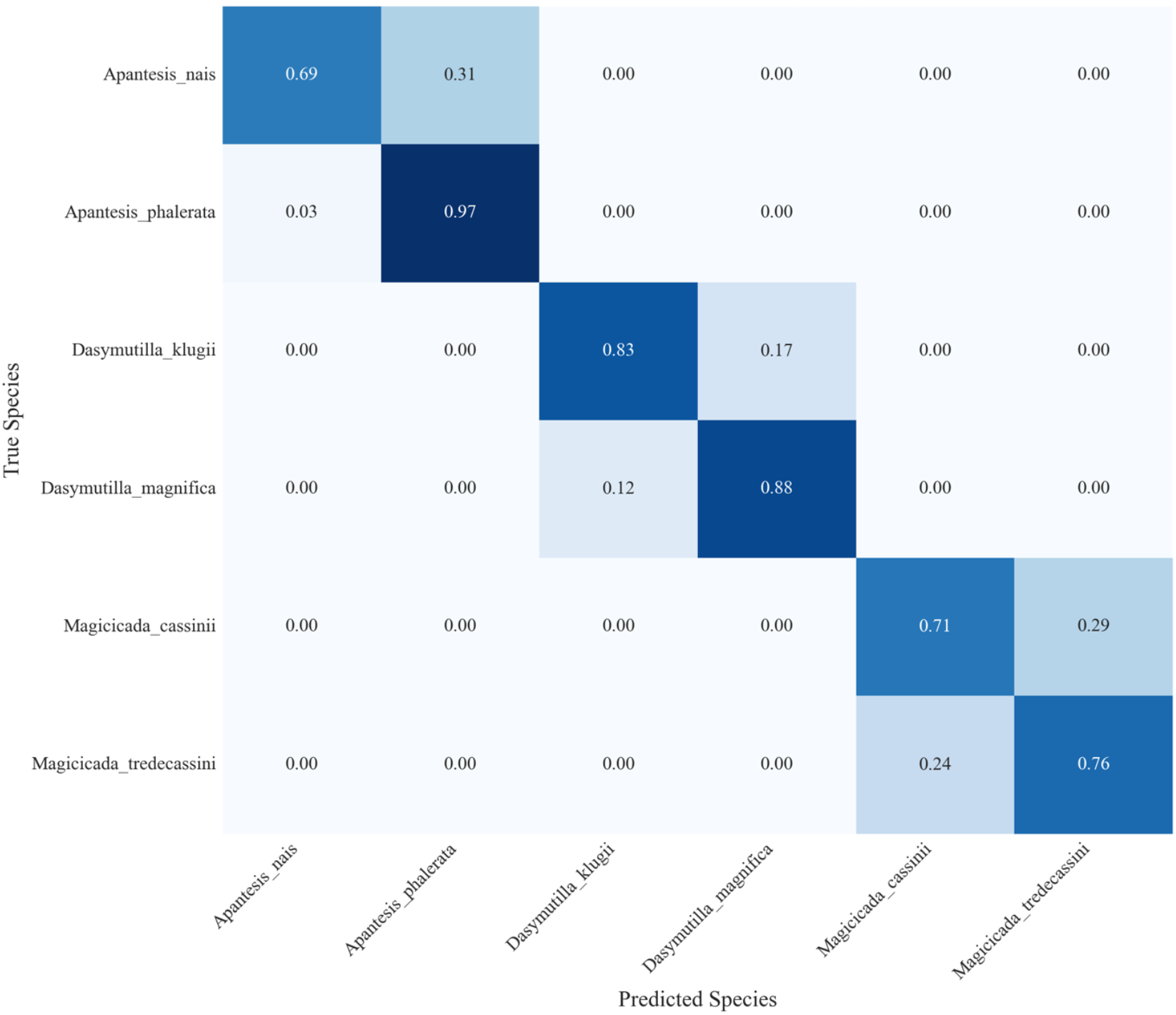
Confusion matrix highlighting cryptic species complexes. Heatmap visualization of the most frequently confused species pairs in the internal test set. Diagonal intensity represents correct classifications; off-diagonal cells indicate misclassifications. Notable confusion clusters include periodical cicadas (*Magicicada spp*.), damselflies (*Dasymutilla* spp.), and morphologically convergent *Apantesis* species. Matrix values represent percentage of predictions; only species pairs with >5% confusion rate are displayed for clarity.

### 3.2 Feature Learning and Interpretability

To validate feature learning, high-dimensional features were visualized using t-SNE (van der Maaten & Hinton, 2008), revealing distinct phylogenetic clustering (Figure 4). Grad-CAM analysis (Selvaraju et al., 2017) confirmed the model attends to diagnostic features such as wing venation and elytral patterns rather than background context (Figure 5).

**Figure 4.**
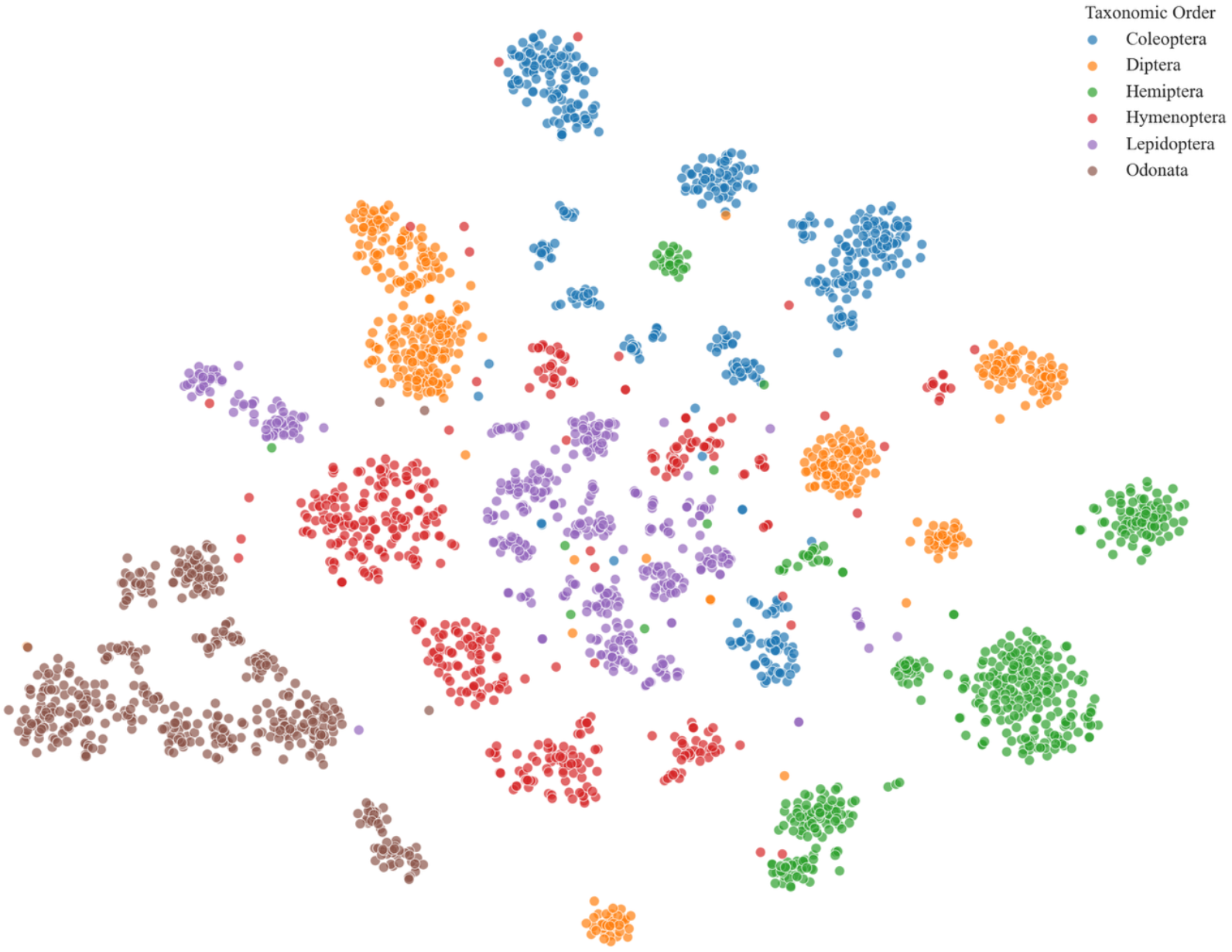
Visualization of the learned feature space using t-Distributed Stochastic Neighbor Embedding (t-SNE). Two-dimensional projection of the 1,280-dimensional feature vectors extracted from the penultimate layer (global average pooling) of the fine-tuned EfficientNet-B0 model. The plot represents a stratified random subset of the internal test set, balanced across six major taxonomic orders. Each point corresponds to a single input image, color-coded by order: Coleoptera (blue), Diptera (orange), Hemiptera (green), Hymenoptera (red), Lepidoptera (purple), and Odonata (brown). The distinct separation between clusters indicates that the model has learned robust, discriminatory morphological representations for high-level taxonomic categories. Furthermore, the visible sub-structures within larger clusters (particularly within Coleoptera and Lepidoptera) suggest the model captures fine-grained hierarchical features corresponding to family- or genus-level similarities without explicit supervision at those levels.

**Figure 5.**
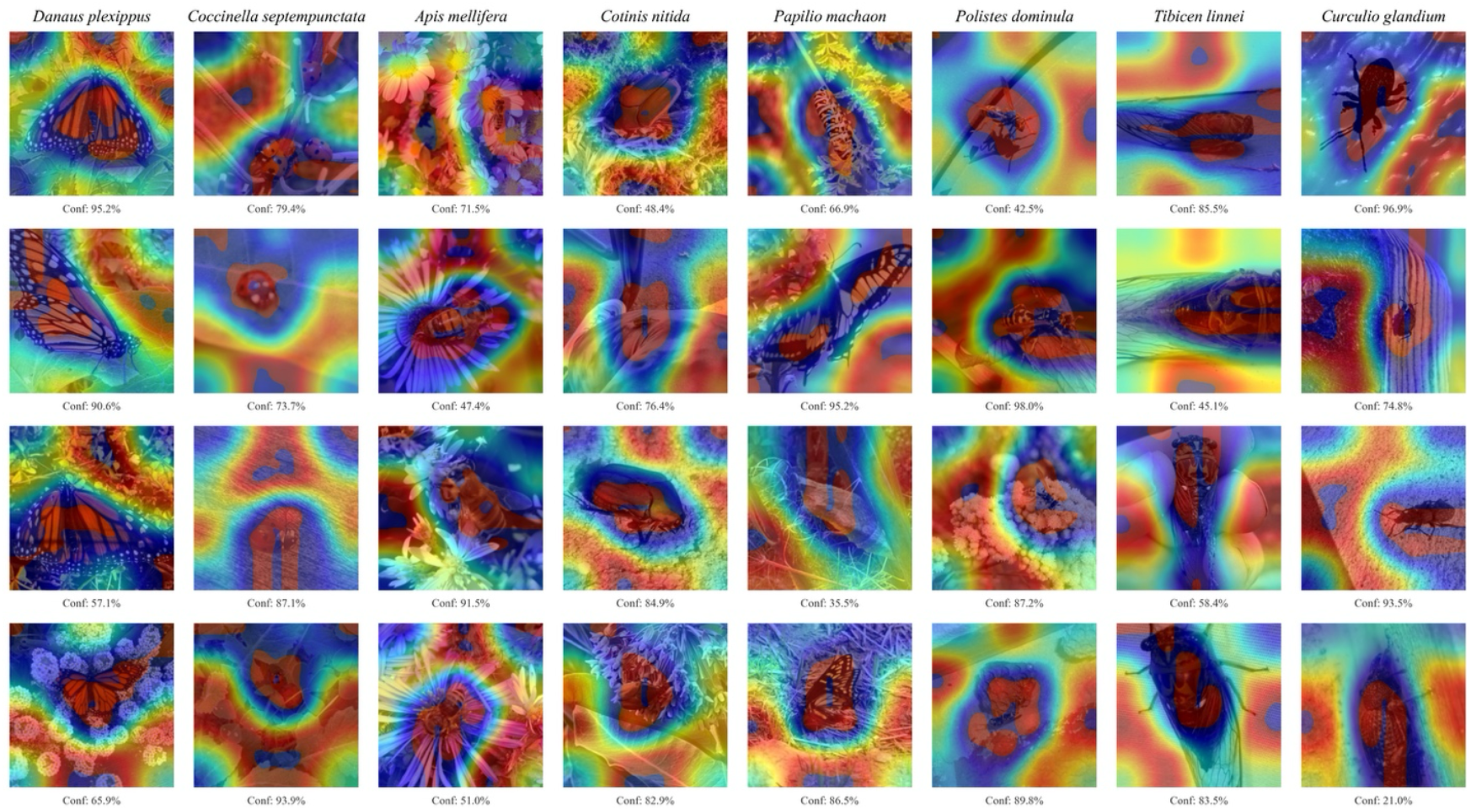
Consistency of feature localization across diverse insect orders using Class Activation Mapping (Grad-CAM). Visualizations represent the gradient-weighted class activation maps derived from the final convolutional layer of the fine-tuned model. a–h, Representative species selected from the internal test set: *Danaus plexippus* (a), *Coccinella septempunctata* (b), *Apis mellifera* (c), *Cotinis nitida* (d), *Papilio machaon* (e), *Polistes dominula* (f), *Tibicen linnei* (g), and *Curculio glandium* (h). Rows display four randomly selected samples (fixed seed n=42) for each species to illustrate model robustness to variations in pose, scale, and background clutter. Color Scale: Warmer colors (red/yellow) indicate regions of high activation that strongly influence the classification decision, while cooler colors (blue) represent negligible contribution. Interpretation: The model demonstrates consistent localization of key morphological features (e.g., wing venation in *D. plexippus*, elytral patterns in *C. septempunctata*) while frequently attending to immediate ecological context (e.g., floral structures associated with *A. mellifera*), suggesting the integration of both organismal and environmental cues for classification. Heatmaps were upsampled via bicubic interpolation and normalized using 10th–99th percentile clipping to suppress background noise.

### 3.3 Inference Efficiency

The final Core ML model size is 30 MB. Inference benchmarks on an iPhone 13 (Apple Neural Engine) yielded a throughput of >700 FPS, confirming suitability for real-time video analysis.

## 4 Discussion

Elytra 1.0 demonstrates that high-accuracy insect identification does not require server-grade transformers. By combining a lightweight architecture (EfficientNet-B0) with rigorous data curation and adaptive training strategies, this model achieves 86.68% observer-independent test accuracy and 91.27% test accuracy while maintaining a 30 MB footprint suitable for offline, edge-native deployment.

The 4.59% generalization gap likely reflects two compounding factors: (1) the strict exclusion of training photographers, and (2) the inadvertent biogeographic shift of the test set. Because the observer-independent test set was collected prospectively during the North American winter (2025–2026), it heavily sampled the southern/Neotropical limits of species’ ranges. This effectively tested the model on a worst-case distribution: identifying temperate species in tropical environments. For example, a Monarch butterfly (*Danaus plexippus*) in the training set is typically photographed on *Asclepias* in a temperate summer meadow, whereas test images likely depicted overwintering individuals in Mexican oyamel fir forests.

Importantly, the model maintained strong performance (86.68%) well above baseline approaches, demonstrating practical utility for real-world deployment where images will originate from diverse, previously unseen contributors. The high photographer diversity in the training data (diversity score: 1.21) likely contributed to this robust generalization. The observer-independent evaluation demonstrates that Elytra 1.0 can serve as a practical tool for biodiversity monitoring in citizen science applications, where photographer diversity is the norm.

Performance variation across taxonomic orders (Figure 2) reflects inherent differences in morphological distinctiveness. High-performing groups like Diptera (92.5%) and Odonata (92.0%) possess diagnostic features, such as wing venation, that are both visible and consistent. Conversely, Hymenoptera’s lower performance (79.1%) stems from widespread cryptic species complexes. For these groups, the 224×224 pixel input resolution likely acts as a primary bottleneck, as diagnostic micro-features are lost during downsampling.

Several methodological choices proved important. The balanced dataset design (median 900 images) prevented model bias toward common species. The high photographer diversity (1.21 observer diversity score) in the training data reduced potential for photographer-specific overfitting, as evidenced by the model’s strong performance on completely novel observers (86.68% test accuracy).

Visual identification reaches an irreducible error floor for cryptic species complexes lacking distinguishing visual features. Periodical cicadas (Magicicada spp.) are primarily differentiated by life cycle timing, information absent from static images (Figure 3). Additional limitations include geographic scope (North American species only), taxonomic breadth (rare species excluded by the observation threshold), and life stage coverage (predominantly adults). While the observer-independent test set is smaller than the full validation corpus, its sample size (N=5,780) provides robust statistical precision (95% CI: ±0.74%) and is comparable to or larger than test sets used in similar fine-grained classification studies (e.g., iNaturalist 2021: N=3,000). The explicit exclusion of training photographers represents a more stringent evaluation than conventional random holdouts, as it directly addresses potential confounding from photographer-specific biases in composition, lighting, and subject presentation.

The achieved test accuracy of 91.27% and observer-independent test accuracy of 86.68% are competitive with modern deep learning approaches for fine-grained species classification, while offering substantial computational and environmental advantages. Elytra 1.0’s lightweight architecture enables edge deployment with significantly lower energy consumption than server-grade models, an important consideration for sustainable, large-scale biodiversity monitoring programs that may deploy thousands of devices continuously for years.

Model training was conducted on a single Apple Mac Studio (M1 Ultra) over 275 hours, consuming approximately 18 kWh of electricity, orders of magnitude less than typical foundation model training runs (Strubell et al., 2019). Training was performed in Burlington, Vermont, where the municipal utility (Burlington Electric Department) provides 100% renewable energy, resulting in effectively zero carbon emissions. This demonstrates that large-scale deep learning on datasets exceeding 2.5 million images can be achieved with minimal environmental impact when paired with efficient hardware and renewable energy grids. The ability to train high-accuracy species classifiers on consumer hardware also democratizes AI development for conservation, enabling researchers in resource-limited settings to develop region-specific models without access to institutional compute clusters.

The 30 MB model size and real-time inference speed make deployment accessible across a wide range of edge devices, from smartphones to low-cost single-board computers. This enables practical applications including autonomous monitoring in remote locations, citizen science validation through mobile applications, and agricultural pest identification for rapid intervention decisions.

Future work could improve cross-photographer generalization through (1) augmentation strategies explicitly targeting photographer-specific biases (e.g., background diversity, lighting variation simulations); (2) photographer diversity requirements during dataset curation; or (3) meta-learning approaches that adapt to novel photographer styles at inference time. Future work will integrate spatiotemporal metadata (location, date, elevation) to resolve geographic variants and bioacoustic data to differentiate cryptic complexes producing species-specific calls. Expanding coverage to larval stages would enhance agricultural utility where immature insects are primary pests.

